# Automated Segmentation of Epithelial Tissue Using Cycle-Consistent Generative Adversarial Networks

**DOI:** 10.1101/311373

**Authors:** Matthias Häring, Jörg Großhans, Fred Wolf, Stephan Eule

## Abstract

A central problem in biomedical imaging is the automated segmentation of images for further quantitative analysis. Recently, fully convolutional neural networks, such as the U-Net, were applied successfully in a variety of segmentation tasks. A downside of this approach is the requirement for a large amount of well-prepared training samples, consisting of image - ground truth mask pairs. Since training data must be created by hand for each experiment, this task can be very costly and time-consuming. Here, we present a segmentation method based on cycle consistent generative adversarial networks, which can be trained even in absence of prepared image - mask pairs. We show that it successfully performs image segmentation tasks on samples with substantial defects and even generalizes well to different tissue types.

## 1 Introduction

Mechanically active tissues, force generating and proliferating cell groups, and topological tissue rearrangement play a critical role for a wide range of biomedical research challenges. Analyzing such dynamical tissues is of central importance to advancing regenerative medicine and stem cell technologies, to understand cancer progression and treatment and for fundamental research into organogenesis and embryo development. Over the past decade, the rapid advancement of photonic imaging technologies has made it relatively easy to collect time lapse imaging data sets of massive size, recording tissue dynamics and rearrangement in vivo over extended periods of time. A prominent bottleneck in the computational analysis of such large-scale imaging datasets is the automated segmentation of cell shapes to quantify tissue topology, geometry and dynamics. Especially the segmentation of images with lower quality remains challenging.

Recently, deep convolutional neural networks were applied successfully in a vast variety of visual recognition tasks, including automatic biomedical image segmentation. In general, their performance is equal or superior to sophisticated rule-based methods [1], [2]. One drawback of these models, however, is the necessity to prepare a well-suited dataset on which the network can be trained. Generating hand-labeled datasets of image - ground truth mask pairs is time-consuming and thus represents an expensive bottleneck. Due to this training bottleneck many practitioners still use rule-based methods instead of machine learning techniques. Moreover, even if extensive training data is available, the performance of these systems degrades significantly when they are applied to test data that differ from the training data, for example, due to variations in experimental protocols. Furthermore, pixel by pixel classifiers based on deep convolutional architectures can perform poorly on image data with substantial defects that have been labeled incompletely due to bleaching and label failure.

Here, we focus on the segmentation of epithelial tissue from *Drosophila Melanogaster* embryos (see figure 1). Drosophila is a common model system due to its simplicity, availability of well explored genetic tools, economic advantages and short life cycles [3]. Furthermore, the size of the organism allows recording of cell tissue in the living, developing embryo. Research in morphogenesis often relies on observations in this model system and in particular the analysis of dynamic processes during morphogenetic phases, like gastrulation, can give important insights about embryonic development [4]. As these processes typically are highly active and involve remodeling and reshaping of cell tissues, a well functioning data acquisition pipeline is crucial to obtain good statistics for quantifying complex dynamics such as coordinated cell oscillations [5], [6], [7]. Generating segmentations of cell tissues in sufficient quantities for such studies is very time-consuming and, depending on the scope of the project, can take months. Therefore, it is highly desirable to automate the segmentation process.

**Figure 1:**
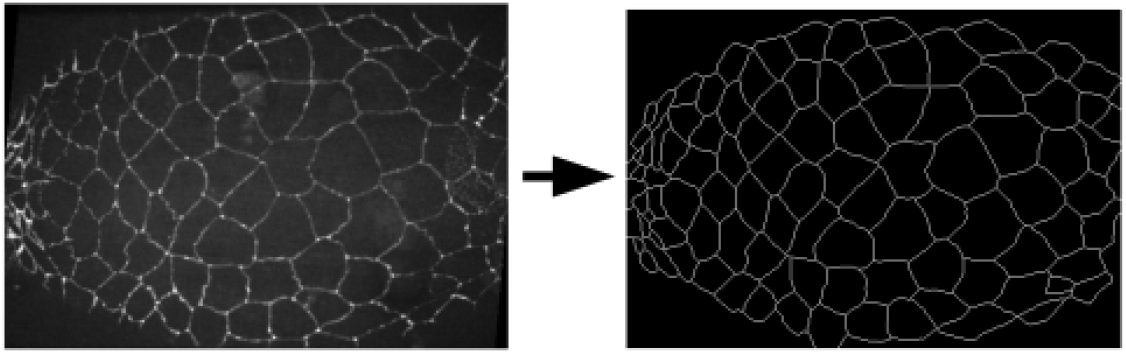
Segmentation of Amnioserosa tissue from Drosophila embryos. For further quantitative analysis a skeletonized version of the microscopy image has to be created such that it contains only cell outlines in a binary representation. From this segmented image, quantities like cell area or cell junction length can be easily extracted.

We propose to employ an automated segmentation pipeline based on cycle-consistent generative adversarial networks (Cycle-GANs [8]). The advantage of this approach is that Cycle-GANs can be trained even in absence of prepared image - mask pairs. We show that this model performs competitively on standard segmentation tasks even when trained on just a few target samples. In addition, our model generalizes well to test data differing from the training data and successfully performs image segmentation tasks on samples with substantial defects. We test this framework on Amnioserosa, an oval shaped tissue that appears at the end of gastrulation in Drosophila morphogenesis.

This paper is structured in the following way: first we briefly review some state of the art methods for cell segmentation, then we introduce our approach based on generative adversarial networks, and finally compare and evaluate the accuracy obtained by our Cycle-GAN approach, a fully convolutional neural network approach (U-Net) and the Tissue Analyzer software, which is commonly used in the field.

## 2 State of the Art Cell Segmentation

We first briefly introduce two state of the art methods, which are commonly used in image segmentation tasks of cell tissues. These methods will then serve as benchmarks for our approach.

### 2.1 Rule-based Methods

To extract cell features from microscopy recordings, practitioners typically employ segmentation methods based on the watershed algorithm, which is e.g. implemented as a plugin in *ImageJ* as the *Tissue Analyzer*[9]. In this plugin, the user can vary the parameters of the algorithm to meet differences in the recording e.g. due to contrast, cell type or bleaching. In addition, segmentation errors can be corrected by redrawing cell contacts by hand, deleting wrongly assigned cell contacts or completely segment the image by hand. If the recording quality is not sufficient for accurate watershed based contour detection, correcting the image by hand can take up to several days for a single movie of 150 frames. The main disadvantage of this method is therefore the time-consuming manual correction of low-quality cell tissue recordings that do not allow for an accurate watershed segmentation. Thanks to the many useful tools for analysis or visualization implemented in the Tissue Analyzer, it is a very popular segmentation program for cell tissues in biology, e.g. [10], [11], [12].

### 2.2 Machine Learning

Machine learning and in particular deep learning approaches are applied in a variety of visual recognition tasks. Especially for biomedical image segmentation they have been shown to be very successful in terms of accuracy and robustness, e.g. [13], [14], [15], [16], [17]. By now there are many competing architectures that yield fast performance and high accuracy. One widely used example is the U-Net, which employs a fully convolutional network architecture with pixel by pixel classification, see [18] for more information. Pixel by pixel classification networks take global information into account but final classification decisions are made based on local information. Therefore, when looking at cell segmentation tasks, errors may happen if recordings are distorted by bleaching effects or gaps in cell contours due to discontinuous labeling. In this study, we take a different approach using so-called *generative networks*, which will be described in the following section.

## 3 Methods

### 3.1 Generative Networks

In contrast to pixel by pixel classifiers, *generative networks* learn general global features of a certain class of images, encoded as a very high dimensional probability distribution [19]. Drawing one sample from this learned distribution corresponds to the creation of a specific image. Since we do not just want to generate a realistically looking segmentation but transform a microscopy recording into its corresponding segmentation (image to image translation) it is necessary to condition the output image on an input. After the network has learned how segmentation images are generated from input images, the network can take a microscopy image as input and generate the corresponding segmentation. The idea of this approach is that the quality of microscopy recordings affects contour detection to a lesser extent since the general shape of target images is already known. Furthermore, we utilize the Cycle-GAN framework as it allows for unpaired training and thus does not require the preparation of dedicated image - ground truth training pairs. Details of these frameworks are described in the following section.

### 3.2 Generative Adversarial Networks

Generative Adversarial Networks [20] (GANs) have proven to be one of the most successful methods in image generation and image manipulation tasks [19]. Originally solely developed for image generation, GANs have recently also been applied to conditional image processing like pix2pix [21], text2image [22], image impainting [23], movies [24] or 3D data analysis [25].

The idea behind GANs is the cunning training procedure, incorporating a so-called *adversarial loss*,meaning that two networks are set up in a game against each other. The first network, called *generator*, is trying to generate realistic images to fool a second network, called *discriminator*, which tries to distinguish between real and fake images. This can mathematically be formulated as a minimax game, given the adversarial loss

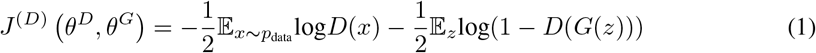

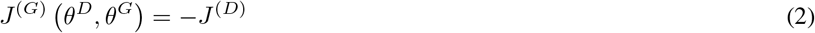

with *J*^(*D*)^ and *J*^(*G*)^ denoting the respective cost functions of discriminator and generator, whereas *θ^D^* and *θ^G^* represent the parameters of both networks to be optimized. Hereby, *D*: *x* → [0, 1] represents the discriminator network, mapping from an image to the probability of this image being real and *G*: *z* → *x* the generator network, whereas *z* is a latent random variable and *x* ~ *p_data_* a sample from the target distribution. Both networks minimize their respective cost function during training. Therefore, the Nash-equilibrium of this game is reached when the discriminator can not decide anymore if the generated image is real or fake, *D*(*G*(*z*)) = 0.5. It can be shown that in this case the probability distribution of the training data is fully recovered [20].

The key aspect is that training happens in a semi-supervised manner, meaning that the discriminator part of the architecture is presented with examples from the generator and the real world and is afterwards informed about its decision being right or wrong. Note that up to this point image to image translation has not entered yet, the generator converts random input into target images, and thus can create images that look like segmentation images but it can not create the corresponding segmentation to a specific tissue recording.

Isola et al. have extended this principle into a unifying framework for image to image translation tasks called conditional generative adversarial networks (cGANs) [21]. Hereby, the generator gets an image as input instead of random variables. Furthermore, decisions of the discriminator now also take the input image into account *D*(*G*(image)|image). At this point, we still need microscopy recording - ground truth segmentation pairs to train the network since the discriminator must be aware of both the correct input and output images. If we omit the necessity of presenting the discriminator with the correct corresponding segmentation it will declare any faithful looking image as being correct. Then it is not guaranteed that the generator learns to create fitting segmentations, it will most likely just convert the input image into random segmentations that look faithful but do not fit the input.

### 3.3 Cycle-GAN

To lift the necessity of microscopy recording - ground truth segmentation pairs in biomedical image processing, we use the Cycle-GAN architecture introduced by Zhu et al. [8]. A simplified sketch of the Cycle-GAN framework is shown in figure 2.

**Figure 2:**
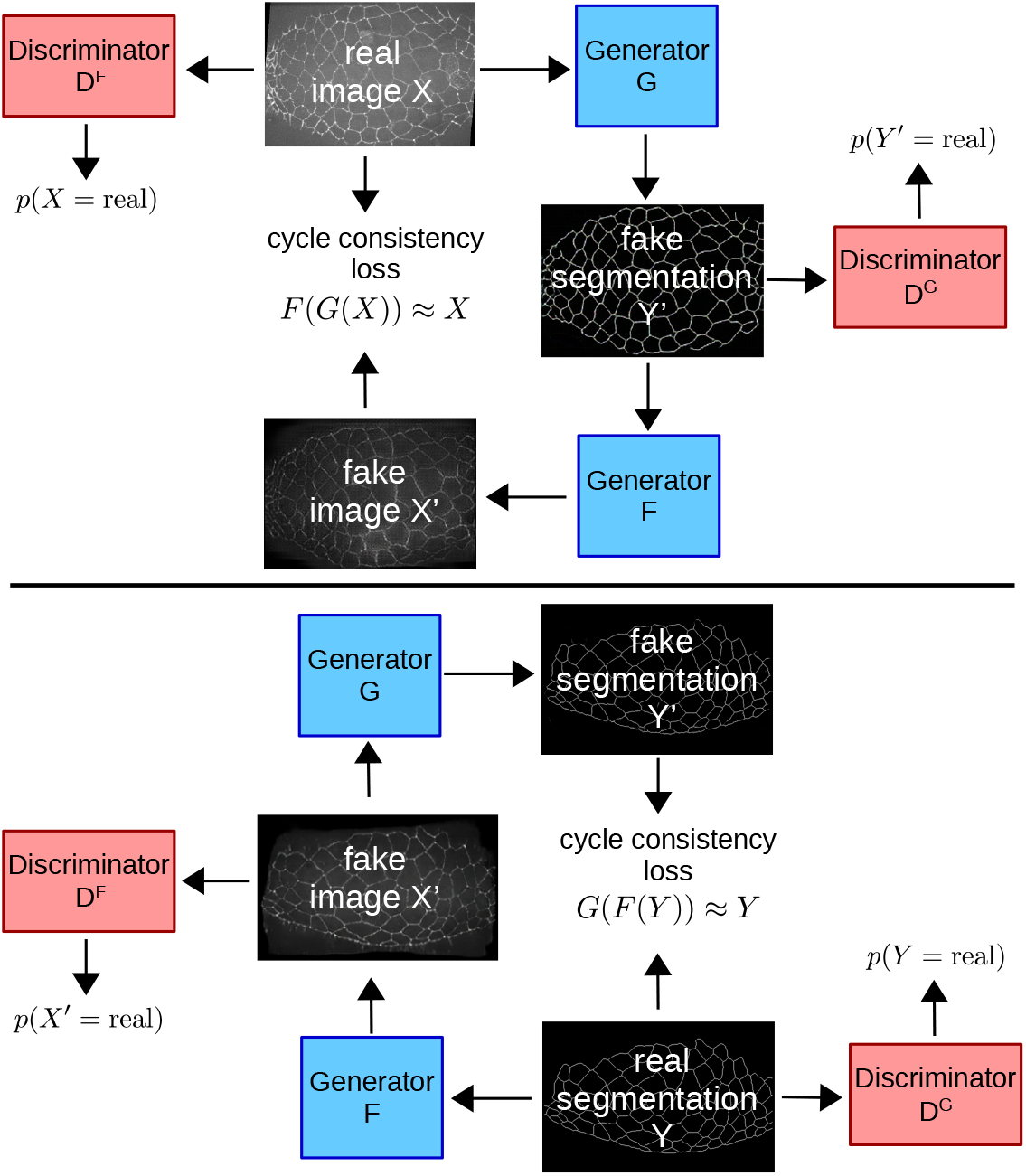
Simplified Cycle-GAN framework. By incorporating a second generator that translates from the segmentation to the microscopy image domain and a cycle-consistent loss that punishes deviations of this outcome from the original microscopy image, the need of dedicated image-ground truth training pairs can be omitted.

The idea behind Cycle-GAN is to enforce a relationship between input domain and target domain by introducing a second generator *F* that translates the generated image of the first generator *G* back into the input domain. Original input image and the target image of the second generator are then compared for similarity. This *cycle-consistency loss* ensures that the target image of the first generator fits to the specific input whereas the adversarial loss ensures that the generator hits the target distribution in the first place.

For the purpose of simplicity we are content with giving an intuitive explanation of the Cycle-GAN framework and refer the interested reader to the literature for a rigorous explanation and further details [8].

We mainly use the original architecture introduced by Zhu et al. with minor changes. For the specific task of image segmentation of epithelial tissues a generator with six Resnet blocks proved to be most successful. We also used a U-Net architecture for the generator with seven down- and upsampling layers, which however showed a slightly worse performance. Although images and and segmentation are grey-scale, we found that the results significantly improved when three colour channels were used. For implementation details we refer to the original paper of Zhu et al. [8].

### 3.4 Training Procedure

The training set for the Cycle-GAN framework consisted of 112 unpaired images of Amnioserosa tissue and ground truth segmentation images, respectively, which were downscaled to a size of 256 × 256 pixels. The images were randomly drawn from a pool of ten, roughly 150 frames long Amnioserosa movies. For evaluation purposes two movies were excluded from the training data: one movie of rather high quality with good contrast and high signal to noise ratio and a second movie with low contrast and discontinuously labeled cell contacts, see figure 3.The low quality movie was picked specifically as a challenging example where watershed might come to its limitations. Depending on the specific setup of the network the training took between 2 - 6 hours on a Titan Xp GPU based on the PyTorch implementation and the hyper-parameters proposed by Zhu et al [8]. After the training, generation of new segmentation images is done at roughly 30fps.

**Figure 3:**
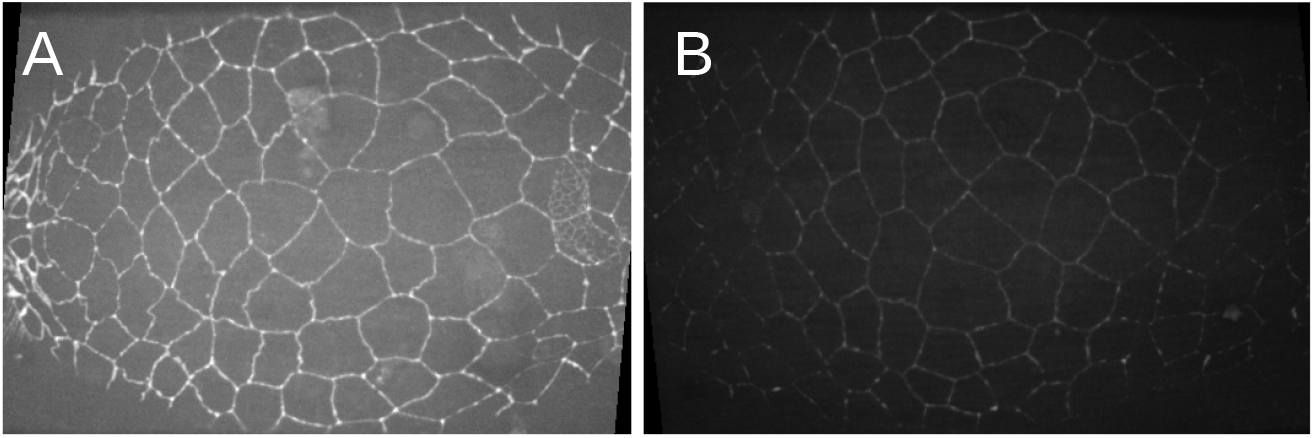
Two movies of Amnioserosa tissue are prepared to test the segmentation accuracy of Cycle-GAN, U-Net and Tissue Analyzer. **A)** Movie of high quality: cell outlines are clearly visible. **B)** Movie of low quality: low contrast makes junctions hard to infer.

### 3.5 Evaluation

Two test sets of Amnioserosa movies were used to compare the performance of three segmentation methods: Tissue Analyzer, U-Net and Cycle-GAN approach.

For the Tissue Analyzer, we choose parameter settings that yield the best possible watershed segmentation. There is no hand correction applied, such that the resulting segmentation will reflect the accuracy of a state of the art watershed algorithm.

For all three methods, the output is a skeletonized version of the input microscopy recording, meaning that the output is a greyscale image with white cell contours and black background. Each method is compared with manually prepared ground truth segmentation images. A fundamental limit to the accuracy tests is therefore the human manual segmentation accuracy.

Segmentation accuracy is measured by the number of correctly identified cells in each frame. Each segmented cell is classified into one of three classes: true positives, cells that are correctly segmented, and false positives, cells that have been segmented although there is no corresponding cell in the ground truth. This can happen if the segmentation method infers additional cell contours for example due to noise or other objects in the recording that are no cell contacts. In addition, cells that appear in the ground truth but not in the segmentation are counted as false negatives.

The comparison between ground truth images and segmentation images of the three methods happens on a pixel by pixel basis. We identify all pixels belonging to a cell in the ground truth and check if these pixels correspond to cells in the output segmentation. Whenever we have an overlap of more than 50% with the ground truth, a cell in the output segmentation is identified as a true positive.

## 4 Results

For the evaluation, we compare the performances of Cycle-GAN, Tissue Analyzer and U-Net. While results show that both machine learning approaches outperform the rule-based approach, it is noteworthy that the accuracy of the Cycle-GAN is at least comparable to the U-Net, which has to be trained on image-mask pairs (figure 4). Even in the challenging case of the lower quality movie, most of the cells are successfully recovered by both machine learning techniques while the watershed method shows a significantly higher failure rate.

**Figure 4:**
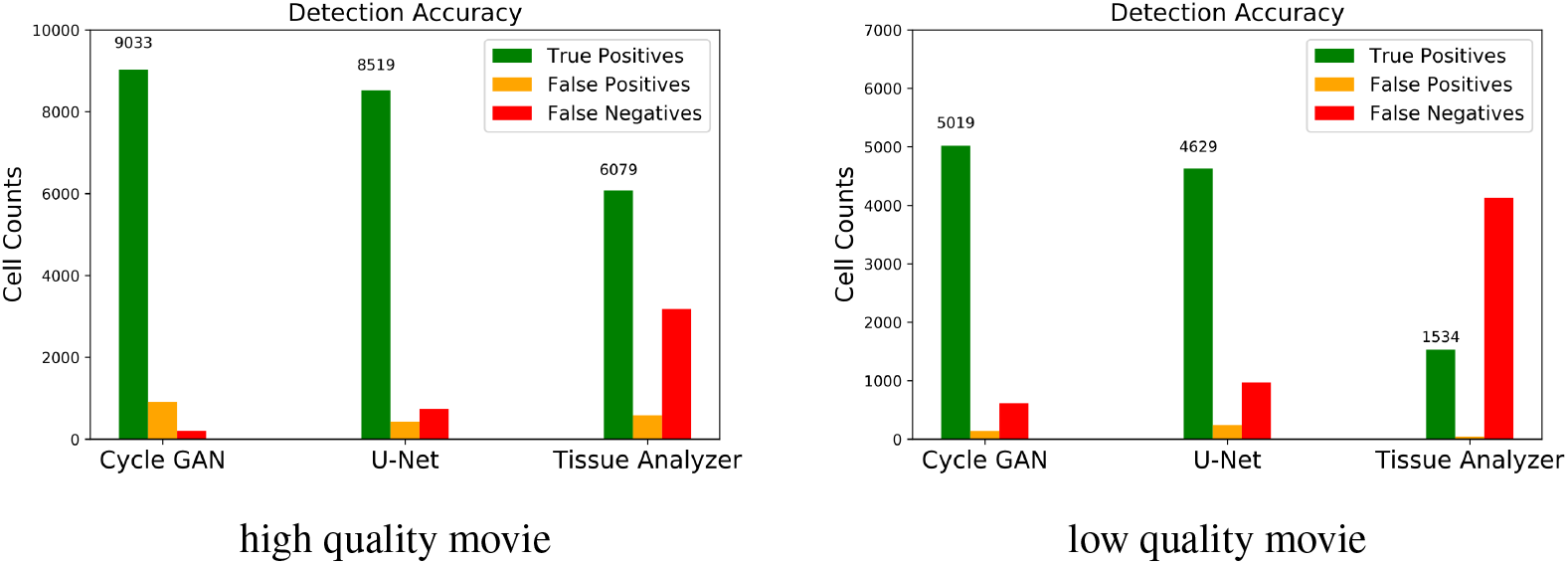
Segmentation accuracy of Cyle-GAN, U-Net and Tissue Analyzer tested with Amnioserosa tissue. Both machine learning methods outperform the rule-based method (without manual correction), whereas Cycle-GAN and U-Net approach achieve comparable accuracy. In the challenging low quality movie case it can be seen that the watershed algorithm reaches its limits, while Cylce-GAN and U-Net correctly detect most of the cells.

From this example we can conclude that the Cycle-GAN framework performs competitively well when compared to the U-Net. It also shows that the Cycle-GAN framework achieves higher accuracies compared to the watershed algorithm.

### 4.1 Domain adaptation

To show the domain adaptation capabilities of the Cycle-GAN framework, we have tested its segmentation performance on different tissue types that are qualitatively different from the tissue on which the network was trained, see figure 5. One of these tissues is a *TMC* mutant, an ion channel knockout, that shows larger cells and often displays yolk granules in the focus plane, which makes segmentation with traditional methods very challenging. The second tissue is called *xit*, presenting elongated cells with wavy cell borders. As a last example, we take tissue from the *germband*, another tissue type that is playing an important role during gastrulation in Drosophila embryos. This tissue is heavily remodeled during the process and therefore more dynamic than Amnioserosa.

**Figure 5:**
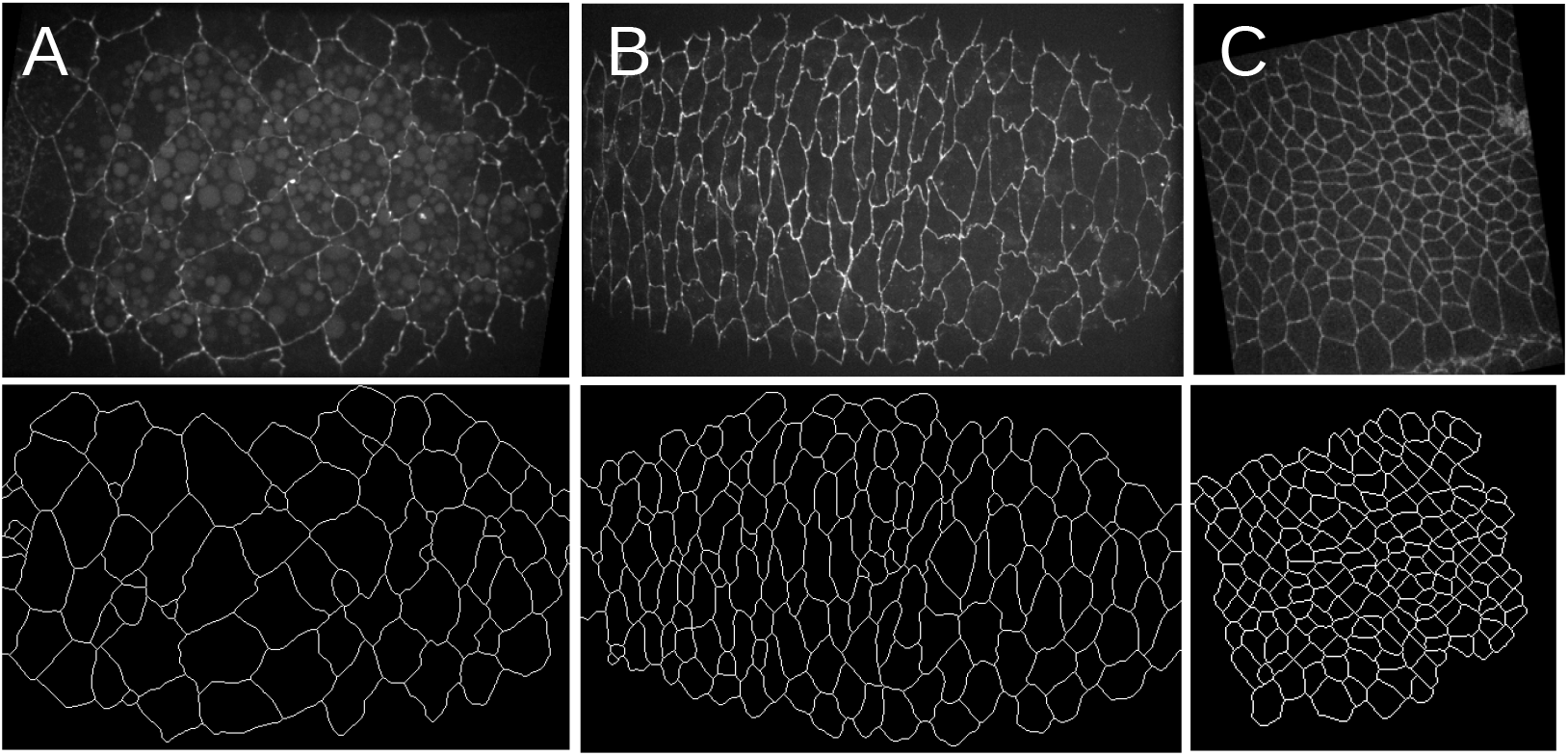
Further examples of Cycle-GAN segmentation in different tissues, trained on the same Amnioserosa data. This comparison shows how well the Cycle-GAN approach is able to generalize from a specific type of training data to similar but qualitatively different tissues. **A)** TMC Amnioserosa mutant: cells are bigger and yolk granules have wandered into focus which makes segmentation very challenging. **B)** Xit Amnioserosa mutant: shows a different phenotype with longer stretched cells and wavy borders. **C)** Epithelial tissue from the germband in Drosophila during gastrulation: continuously reshaping tissue type that consists of smaller cells.

We can see that the overall performance is still very good, but at the same time limitations of this method become clear. In the case of the TMC mutant, it is impressive how accurate the segmentation is despite the original image being highly distorted by yolk granules that moved into focus. There are however a number of wrongly assigned cell contacts where the network identified the outline of a yolk granule as cell outline. The Xit mutant is accurately segmented as well, but the network fails to show accuracy on small scale details of the cell junctions, in particular the wavyness of some cell borders is lost. This effect is due to the training on Amnioserosa wild type tissue where cell contacts are exclusively smooth and thus while being able to identify the overall cell shapes the microscopic structure is lost. Interestingly, in the third case of germband tissue we obtain very high accuracy without visible errors. The main difference to the Amnioserosa tissue is that cells are much smaller and more dynamic in time, whereas the overall shape of the cells is similar.

These examples show that the Cycle-GAN framework is very robust and shows high performance on the specific tasks it was trained for. It can however not adapt to identify qualitatively different shapes as seen in the case of wavy cell borders. On the other hand, if the general shape of cells is similar to the training data, like in the TMC case, then high accuracy can be achieved despite distortion of the image by yolk granules.

## 5 Conclusions

In this paper we showed that the problem of ground truth creation for training of deep convolutional networks can be circumvented using a Cycle-GAN framework. Our results indicate comparable performance to other deep learning methods and advantages over traditional rule-based methods. In addition, Cycle-GANs shows high potential on cell tissues that differ from the training set.

Future work has to explore generalization to different segmentation tasks as this study is limited to the segmentation of epithelial cell tissue.

